# Validation of SNP markers for marker-assisted selection of genotypes with increased carotenoid and dry matter contents in cassava

**DOI:** 10.1101/2022.01.27.478005

**Authors:** Adenike D. Ige, Bunmi Olasanmi, Guillaume J. Bauchet, Ismail S. Kayondo, Edwige Gaby Nkouaya Mbanjo, Ruth Uwugiaren, Sharon Motomura-Wages, Joanna Norton, Chiedozie Egesi, Elizabeth Y. Parkes, Peter Kulakow, Hernan Ceballos, Ismail Y. Rabbi

## Abstract

Provitamin A biofortification and increased dry matter content are important breeding targets in cassava improvement programs worldwide. Biofortified varieties contribute to the alleviation of provitamin A deficiency, a leading cause of preventable blindness in developing countries. Dry matter content is a major component of dry yield and thus underlies overall variety performance and acceptability by growers, processors, and consumers. SNP markers linked to these traits have recently been discovered through several genome-wide association studies but have not been deployed for marker-assisted selection (MAS). Assessment of marker performance in diverse genetic backgrounds is an important step towards their deployment for routine MAS. In the present study, seven previously identified markers for these traits were converted to a robust set of uniplex allele-specific PCR assays and validated in two independent pre-breeding and breeding populations. These assays were efficient in discriminating marker genotypic classes and had an average call rate greater than 98%. A high correlation was observed between the predicted and observed carotenoid content as inferred by root yellowness intensity in the breeding (r = 0.92) and pre-breeding (r = 0.95) populations. On the other hand, dry matter content-markers had moderately low predictive accuracy in both populations (r < 0.40) due to the more quantitative nature of the trait. This work confirmed marker effectiveness in multiple backgrounds, therefore, further strengthening its value in cassava biofortification to ensure nutritional security as well as dry matter content productivity. Our study provides a framework to guide future marker development, thus leading to more routine use of markers in MAS in cassava improvement programs.

## 1.0 Introduction

Cassava (*Manihot esculenta* Crantz) is a principal starchy root crop for both the rural and urban populations in the tropics and sub-Saharan Africa in particular. The continent accounts for more than half of the total world’s production of 304 million tonnes (FAOSTAT, 2020). Due to its ability to grow with little agricultural inputs in marginal environments characterized by poor soils and water stress, the crop takes on the crucial role of being a key food security crop in sub-Saharan Africa (Burns et al., 2010).

Among the major staple sources of carbohydrates, cassava has one of the longest breeding cycles ranging from five to eight years (Ceballos et al., 2004, 2012). This is due to its long growth cycle of 12 - 18 months; clonal propagation which results in low multiplication rates of planting propagules; its high levels of heterozygosity; and difficulty in making crosses due to poor and asynchronous flowering as well as low seed set per cross (Jennings and Iglesias, 2002; Ceballos et al., 2012). These challenges notwithstanding, breeding programs around the world have developed improved varieties that address various production constraints including biotic and abiotic stresses, improved yield and dry matter content (Kawano 2003; Okechukwu and Dixon 2008) as well as enhanced micronutrient content, particularly of provitamin A carotenoid (Andersson et al., 2017; Ilona et al., 2017). However, as the demand for cassava for food, feed, and industrial raw material continues to grow due to increase in population (Anyanwu et al., 2015; Parmar et al., 2017), breeding programs need to adopt modern breeding technologies and tools such as marker-assisted selection or genomic selection in order to increase the rate of genetic gain to meet the demands in an ecologically sustainable manner (Ceballos et al., 2015).

Marker Assisted Selection (MAS) is one of the most important applications of molecular marker technology in plant breeding (Collard and Mackill, 2008). It facilitates the indirect selection of new plants based on the presence of a favorable allele at a marker that is closely linked to a trait of interest (Collard and Mackill, 2008). In cassava, MAS can be used at the early stages of the breeding scheme to select individuals with favorable alleles for storage-root traits that would otherwise only be phenotypically evaluated at maturity. This has several advantages, namely: 1) reduction in the time it takes to make a decision to advance a clone to the next stage of testing; 2) reduction in the number of clones to be advanced to larger plot trials thereby saving scarce phenotyping resources, and 3) in some cases, the cost of marker assay is lower than those that are usually expended on the actual trait phenotyping. A good example is carotenoids quantification using spectrophotometry method and High-performance Liquid Chromatography (HPLC) which can be many-fold more expensive than a SNP assay (Semagn et al., 2014; Andersson et al., 2017). Therefore, the adoption of MAS can increase the efficiency of selection, leading to a more rapid rate of genetic gain, fewer cycles of phenotypic evaluation thus, reducing the time for varietal development (Collard and Mackill, 2008).

The prerequisite for the application of MAS is the identification of major genes or genomic regions associated with a trait of interest. Over the last 15 years, quantitative trait locus (QTL)- mapping studies of different traits in cassava have been published (Akano et al., 2002; Balyejusa et al., 2007; Fregene et al., 2001; Morillo et al., 2013; Okogbenin et al., 2012; Rabbi et al., 2014). Most of these studies used segregating populations developed from either selfed or bi-parental crosses of parents with contrasting trait levels (Rabbi et al., 2014). More recently, association or linkage disequilibrium mapping using a genome-wide association study (GWAS) became an approach for unraveling the molecular genetic basis underlying the natural phenotypic variation (Davey et al., 2011). The advantage of GWAS over QTL mapping is the higher mapping resolution and the identification of a broader set of alleles in large diverse germplasm (Yu and Buckler, 2006). Several GWAS have been conducted on key cassava traits, including CMD resistance (Rabbi et al., 2020; Wolfe et al., 2016), carotenoids content (Esuma et al., 2016; Ikeogu et al., 2019; Rabbi et al., 2017, 2020) and dry matter content (Rabbi et al., 2020) in diverse cassava populations to discover significant loci.

Despite this progress, the output from discovery research has not been translated into assays that breeders can easily use to support selection decisions (Chagné et al., 2019). To overcome this bottleneck and bridge the gap between discovery and routine usage, new trait-linked markers must be technically and biologically validated, preferably using independent populations (Platten et al., 2019; Ige et al., 2021). This process informs the breeder whether the expected allelic phenotypic effects are reproducible in different genetic backgrounds from the one in which the marker-trait association was originally identified (Li et al., 2013).

In this study, we describe the conversion and validation of SNP markers associated with increased provitamin A carotenoid biofortification and dry matter content; two important traits under active improvements in many breeding programs in the world. Although cassava is very efficient in carbohydrate production, its starchy roots lack essential micronutrients including provitamin A carotenoid (Sayre et al., 2011; Ceballos et al., 2017). Vitamin A deficiency (VAD) often leads to several severe health and economic consequences including increased incidence of night blindness, suppressed immunity leading to increased mortality rate especially among pregnant women and young children as well as reduced productivity (Sayre et al., 2011; WHO, 2020). Dry matter content is a crucial yield component and is a key determinant of variety acceptance by growers, processors, and consumers (Sánchez et al., 2014; Bechoff et al., 2018). Varieties with low dry matter content (less than 30%) are often less preferred than those with moderate to high dry matter. Like carotenoid content, dry matter content can only be assessed on mature storage roots at the end of the growing season. Marker-assisted selection is expected to provide breeders with the ability, for example, to screen either for genotypes with high levels of these traits or eliminate those with undesirable levels at early stages of testing, thereby allocating their limited field plots to high-value genotypes.

## 2.0 Materials and methods

### 2.1 Retrieving significant loci linked to increased carotenoid and dry matter contents

Marker discovery, development, and validation workflow used in the present study is presented in Figure 1. The SNP markers linked to increased carotenoid and dry matter contents validated in the present study (Table 1) were derived from Udoh et al. (2017) and Rabbi et al. (2020). Sequencing of four carotenoid pathway candidate genes in 167 cassava accessions from the International Institute of Tropical Agriculture (IITA), Nigeria, uncovered two important SNPs on phytoene synthase 2 (PSY2) (Udoh et al., 2017) The most significant SNP on PSY2 (position 572) is a causal mutation resulting in a non-synonymous amino acid substitution (Welsch et al., 2010). This marker was converted to a KASP assay and renamed as per its chromosomal position on the version 6.1 reference genome (S1_24155522). Additional markers associated with the study traits were obtained from a recent GWAS using a large panel of 5130 diverse clones developed at IITA in Nigeria (Rabbi et al., 2020). The population was genotyped at more than 100K genome-wide SNP markers via genotyping-by-sequencing (GBS). For carotenoid content, a major locus on chromosome 1 tagged by three markers (S1_24159583, S1_24636113, and S1_30543962) as well as five new genomic regions associated with this trait on chromosomes 5, 8, 15, and 16 were identified. Of these, three (S1_30543963, S5_3387558, and S8_25598183) were selected for KASP conversion and validation in the present study. The markers associated with dry matter were S1_24197219, S6_20589894, and S12_5524524.

**Figure 1:**
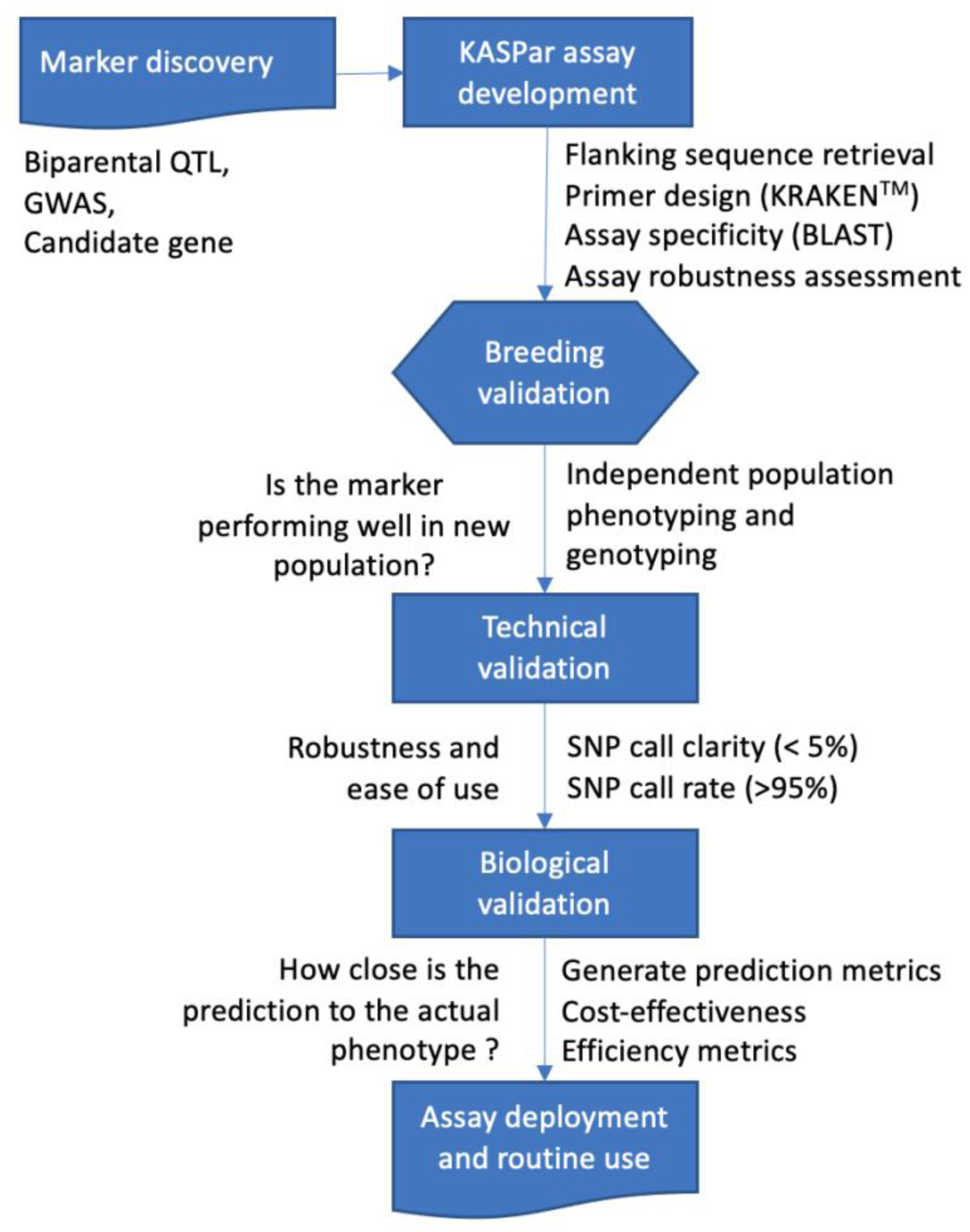
Schematic overview of marker discovery, assay development, and validation of trait-linked markers for molecular breeding

**Table 1:**
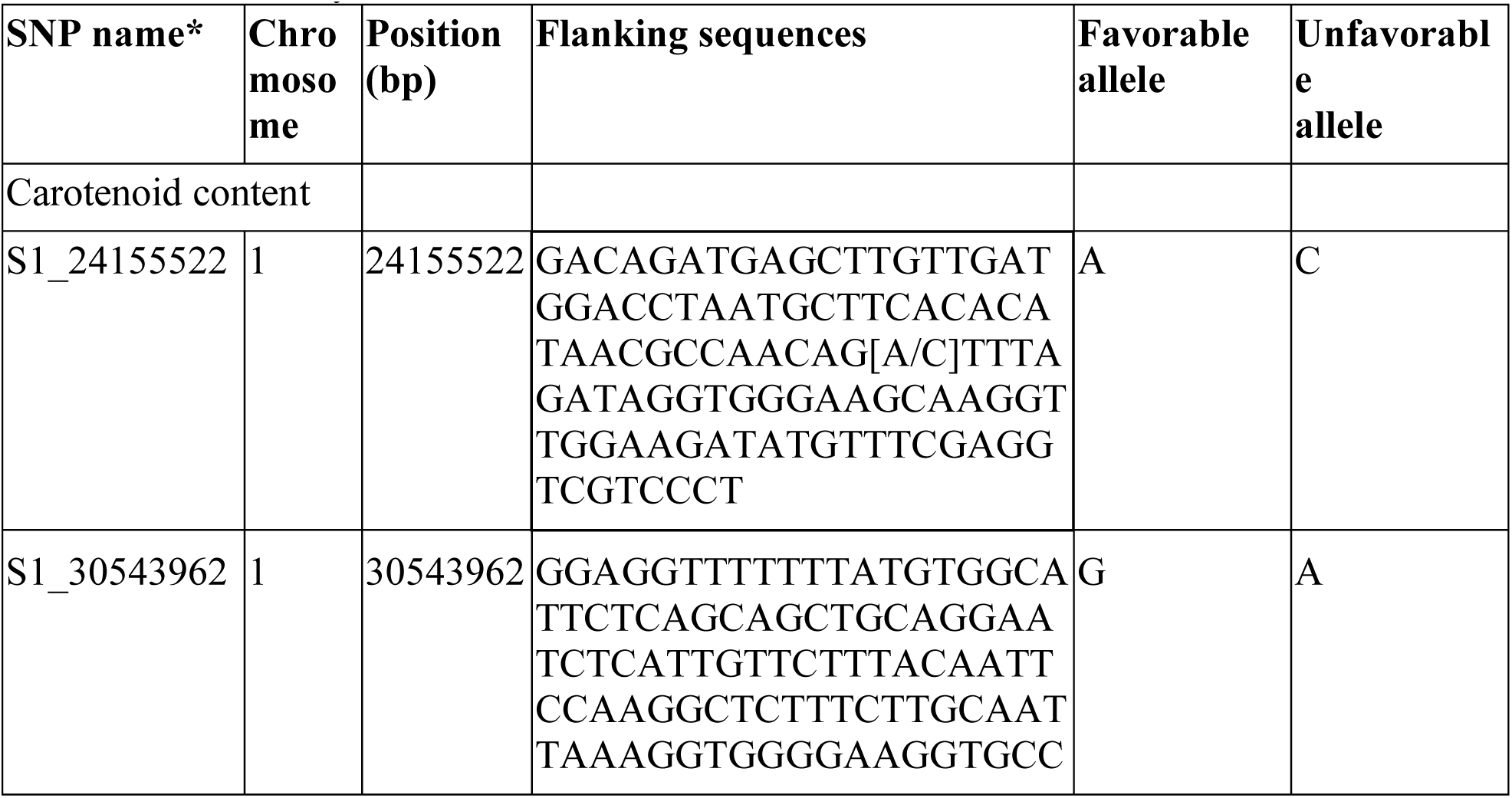
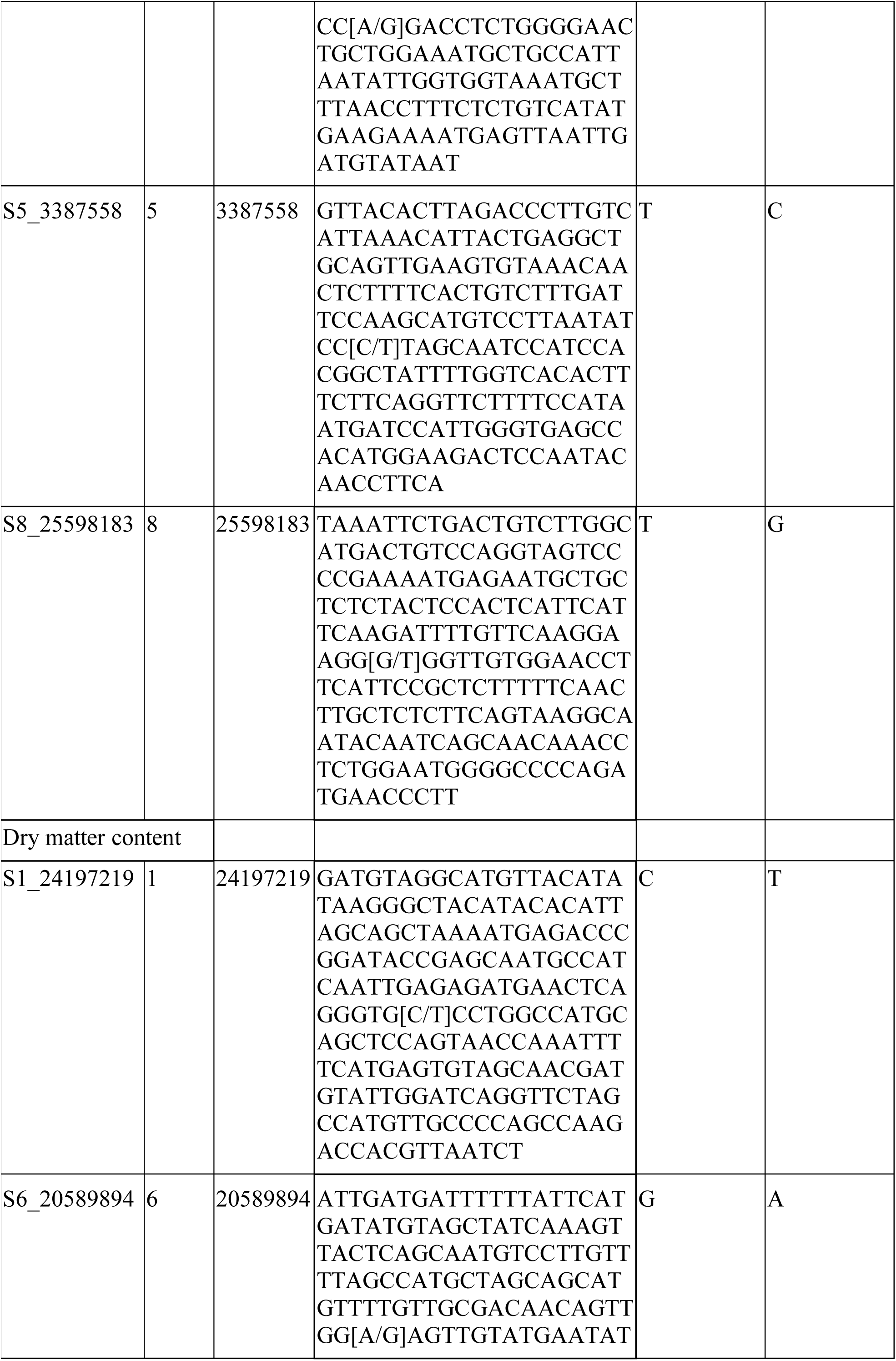
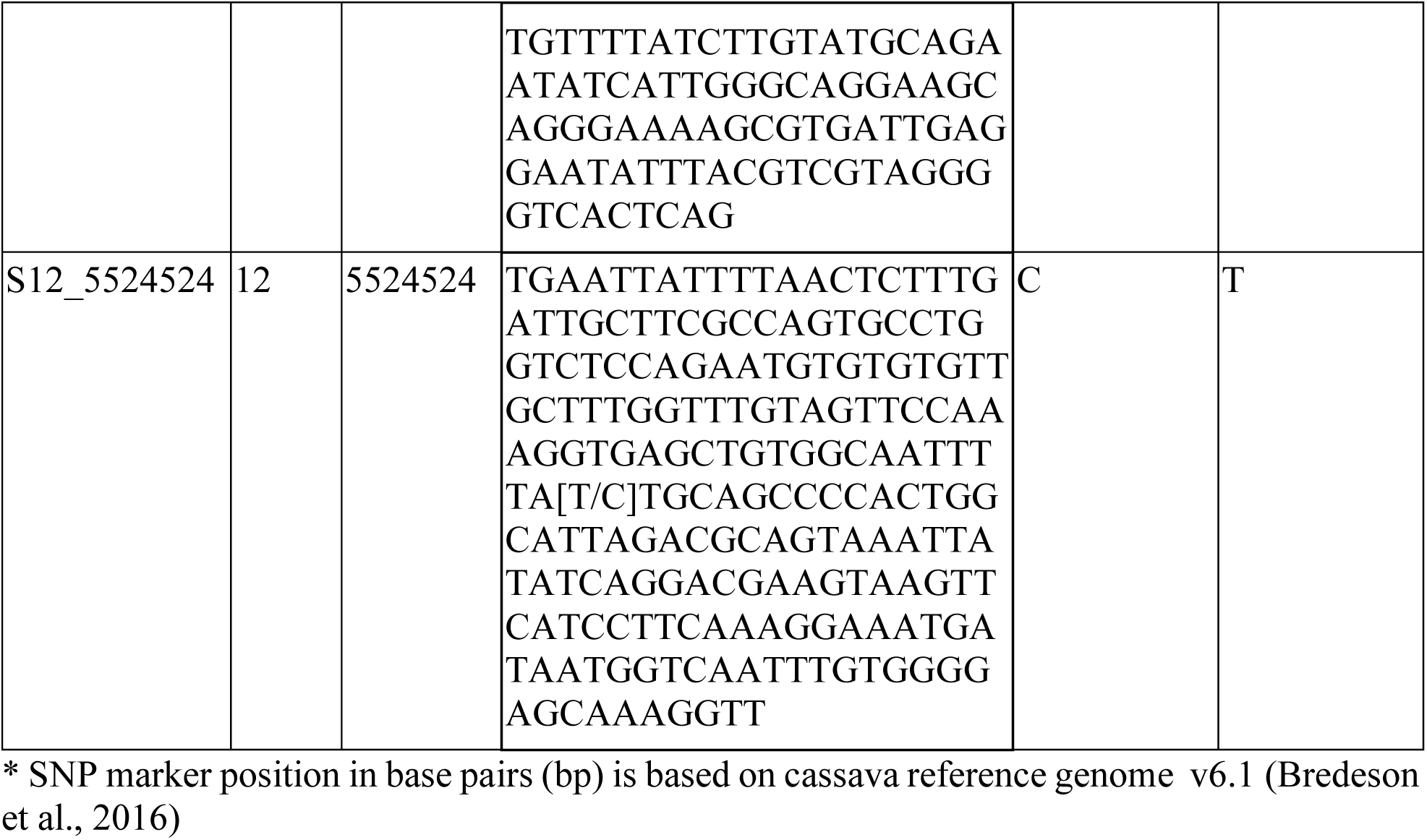
List and description of the seven SNP markers converted to KASP assays and validated in this study

### 2.2 Development of KASP assays

Fifty nucleotide bases flanking the target SNP on each side were obtained from the cassava (*Manihot esculenta*) reference genome (version 6.1) available at https://phytozome-next.jgi.doe.gov/info/Mesculenta_v6_1. Then, a nucleotide-nucleotide Basic Local Alignment Search Tool (BLAST) was used to check for locus-specificity of the assays in order to minimize the possibility of cross-amplification of the marker in non-target regions of the genome. Primers were designed using a proprietary Kraken™ software system from LGC Biosearch Technologies, UK, with the default parameters.

Assay technical validation was carried out using a panel of about 200 genetically diverse cassava accessions that are known to segregate at the SNP assays. A no-template control was included in the SNP genotyping. The robustness of the assays was assessed under four DNA concentrations (Dilution 1= 10X, Dilution 2 = 100X, Dilution 3 = 24X, Dilution 4 = 240X) using metrics such as ease of scoring the three expected genotype classes, tightness, and distinctiveness of the genotypic classes on cluster plots, percentage call rate, and percentage clarity.

### 2.3 Validation of KASP assays in independent populations

The KASP assays’ performances were assessed in two independent populations from IITA, Nigeria. These populations consisting of breeding and pre-breeding germplasm were different from the GWAS marker discovery panel.

#### 2.3.1 Description of the study populations

The breeding population is part of IITA’s regular recurrent selection pipeline and has been derived from controlled crosses among elite genotypes carried out in 2017. Yield, multiple stress tolerance, and dry matter content are the major traits for improvement in this population. The cohort was evaluated initially at the seedling nursery (SN) stage consisting of 22,420 progenies from 563 families (mean family size of 40, ranging from 1 to 220) in 2018 in Ibadan, Nigeria (7°24′ N, 3°54′ E; 200 m above sea level). The SN trial was planted at a spacing of 1 m × 0.25 m and harvested 12 months after planting (MAP); a selection of 1590 genotypes based on disease resistance, plant vigor, plant architecture, and root yield was advanced to clonal evaluation trial (CET) at Ikenne, Nigeria (6°52′ N 3°42′ E; 61 m above sea level).

The pre-breeding population was developed using a polycross hybridization between twenty-three (23) IITA and nineteen (19) CIAT (International Center for Tropical Agriculture) parental clones. To ensure safe germplasm exchange between Africa and Latin America, the hybridization was carried out in Hawaii which has a mild tropical climate that is suitable for cassava survival as well as prolific flowering. The objective of developing the population was to enhance provitamin A biofortification by introgression of a new source of novel alleles for Africa and to develop germplasm incorporating resistance to cassava mosaic disease, high content of provitamin A and starch, and tolerance to acid soils and drought for Latin America. Like the breeding population, a SN evaluation trial was established in Ibadan for 5,608 genotypes planted at a spacing of 1m x 0.25m. The mean family size was 16, ranging from 1 to 165 clones. The trial was harvested 10 MAP and about 7% of the genotypes (790) were advanced to CET at Ikenne, Nigeria (6°52′ N 3°42′ E; 61 m above sea level) based on vigor alone.

#### 2.3.2 Field trial design and phenotyping of cassava storage roots for carotenoid content and dry matter content

Genotypes at the first CET were used for the validation study. Clonal evaluation trial was preferred because of the large size (typically several hundred) and diversity for most of the traits. The only selection imposed in the SN was resistance to diseases, plant architecture as well as poor root yield. The trials were laid out in an augmented design to accommodate the large number of entries at the CET. Each genotype was planted at a spacing of 1m between rows and 0.5m within rows. For the breeding population, 58 plots per 30 sub-blocks with five checks (IITA-TMS-IBA00070, IITA-TMS-IBA30572, TMEB419, IITA-TMS-IBA982101, IITA-TMS-IBA980581) randomly assigned to each sub-block. This trial was planted and harvested in June 2018 and June 2019, respectively. The pre-breeding population’s trial carried out between October 2018 and October 2019 consisted of 900 plots (50 plots per 18 sub-blocks) with four checks (TMEB419, IITA-TMS-IBA30572, IITA-TMS-IBA070593, and IITA-TMS-IBA000070) in each block. All field management practices were performed according to the technical recommendations and standard agricultural practices for cassava (Abass et al., 2014; Atser et al., 2017).

Direct estimation of total carotenoid content using laboratory extraction followed by spectrophotometry and HPLC is not only expensive but also low throughput for routine germplasm screening particularly at the early stages of breeding selection. Due to the large number of accessions in this study, we used two color-based methods to assess the relative difference between cassava genotypes in carotenoid content. Utilization of color intensity as a proxy for the carotenoids content in cassava is justified because of the well-established linear relationship between root yellowness and total carotenoids content (Pearson’s coefficient, r, ranges from 0.81 to 0.84) (Iglesias et al., 1997; Chávez et al., 2005; Marín Colorado et al., 2009; Sánchez et al., 2014; Esuma et al., 2016) as well as with total beta-carotene (Udoh et al., 2017). Moreover, 80 to 90% of total carotenoid content in cassava is provitamin A compared to other crops, making color-based assessment a good proxy for estimating not only total carotenoids content but also total β-carotene content (Wong et al., 2004; Ceballos et al., 2017). In maize, kernel color is not correlated with the primary carotenoid of interest, that is, β-carotene which has the highest pro-vitamin A activity due to the presence of other carotenoids such as β-cryptoxanthin, zeaxanthin, and lutein (Wong et al., 2004)

The first method is a standard visual assessment of root yellowness using a color-chart with a scale ranging from 1 (white root) to 6 (deep yellow root) (Supplementary Figure 1). The second method is a surface color measurement using a CR-410 chromameter (Konica Minolta). The chromameter’s three-dimensional color space defined by L*, a*, and b* coordinates provides a more objective and precise assessment of surface color and its intensity. The Commission Internationale de l’Éclairage (CIELAB) L* coordinate value represents sample lightness ranging from 0 (black) to 100 (diffuse white). The a* values represent red (positive coordinate values) or green (negative coordinate values). Of importance in our study is the b* coordinate whose positive values measure the degree of yellowness and therefore an indirect estimate of carotenoid content.

For the chromameter color measurements, eight roots per plot were peeled, washed, grated, and thoroughly mixed. A subsample was transferred into a transparent sampling bag (Whirl-Pak™) and scanned at four independent positions. The instrument was calibrated each day using a white ceramic and illuminant D65 was used as a source of light.

Root dry matter contents were assessed using the oven-drying method. Eight fully developed roots were randomly selected from each plot, peeled, washed, grated, and thoroughly mixed. For each sample, 100 g was weighed and oven-dried for 72 h at 80°C. The dry samples were then weighed, and the dry matter content was expressed as the percentage of dry weight relative to fresh weight.

#### 2.3.3 Genotyping

Young leaves were sampled at three MAP from the evaluation plots. Three 6mm diameter leaf discs were obtained from each genotype into 96-well plates on ice, and freeze-dried for at least 72 hrs. The samples were shipped to a genotyping service provider (Intertek, Sweden) for automated DNA extraction and SNP genotyping using four (4) markers linked to increased carotenoid content and three (3) markers linked to increased dry matter content (Table 1) using KASP assay. Two blank controls were included in each plate during genotyping.

The KASP assay protocol is provided in the KASP manual (LGC, 2013). In brief, genotyping was carried out using the high-throughput PCR SNPline workflow using 1 μL reaction volume in 1536-well PCR plates. The KASP genotyping reaction mix comprises three components: (i) sample DNA (10 ng); (ii) marker assay mix consisting of target-specific primers; and (iii) KASP-TF™ Master Mix containing two universal FRET (fluorescence resonant energy transfer) cassettes (FAM and HEX), passive reference dye (ROX™), Taq polymerase, free nucleotides, and MgCl_2_ in an optimized buffer solution. The SNP assay mix is specific to each marker and consists of two Kompetitive allele-specific forward primers and one common reverse primer. After PCR, the plates are fluorescently read, and allele calls are made using KRAKEN™ software.

#### 2.3.4 Data analysis

##### 2.3.4.1 Phenotypic data analysis

A linear mixed model was used to obtain the best linear unbiased predictions (BLUPs) for each genotype in the CETs of breeding and pre-breeding populations. The model was fitted using the *lme4* package (Bates et al., 2015) in R software version 4.0.3 (R Development Core Team, 2020). Checks were considered as fixed effects while genotypes and blocks were considered as random effects. The mathematical model used for the incomplete block design analysis is represented as follows:

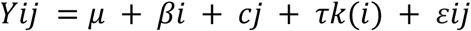

where Y_ij_ is the vector of phenotype data, μ is the grand mean, β is the block effect, c_j_ is the check effect, τ_k(i)_ is the genotype effect, and ε_ij_ is the residual term. Broad-sense heritability was calculated as:

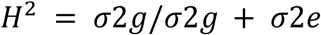

where H^2^ is the broad-sense heritability; σ^2^_g_ and σ^2^_e_ are the variance components for the genotype effect and the residual error.

Pairwise correlation analysis of the traits using the BLUP estimates was determined using the *corr*.*test* function in the psych package (R Development Core Team, 2020).

### 2.3.4.2 Technical and biological validation of KASP markers

Technical performance metrics used to validate the robustness of markers include SNP call rate and call clarity. Call rate is the proportion of samples with non-missing genotype calls. Call clarity is defined by the ease of assigning samples to a genotype class based on their position on a fluorescence cluster Cartesian plot. The tighter and more distinct the cluster, the easier and consistent it is to call the respective genotype class, namely homozygous for either allele 1 or 2 or heterozygous in the case of biallelic SNPs and a diploid genome.

Biological validation of the converted markers was assessed using three complementary approaches. First, the allele substitution effect was visualized using boxplots, and the difference in carotenoid and dry matter content BLUP values among the genotypic classes at each marker locus was assessed using a pairwise t-test. Second, the predictive ability of the SNP markers was estimated using a multiple linear regression model. Marker alleles and the observed phenotypes were considered as the independent and response variables, respectively, as shown in the linear model below:

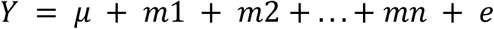

where: *Y*= phenotypic observations of traits, μ = overall mean of the population, m_1_, m_2_, … m_n_ = marker effects, e = residual value.

Bootstrap resampling was carried out to obtain robust estimates of model parameters, specifically the magnitude and confidence intervals of the allele-substitution effects for the markers associated with the two traits (Davison and Hinkley, 1997). The *reg_intervals* function in the tidymodels R package (Kuhn and Wickham, 2020) was used to generate 1000 bootstrap resamples and fit the multiple linear regression model on each one.

Finally, a 5-fold cross-validation analysis repeated 10 times was carried out to obtain marker performance metrics including predictive accuracy (R^2^), root mean square error (RMSE, the square root of the mean squared difference between observed and predicted trait values), and mean absolute error (MAE, the average absolute difference between the predictions made by the model and the actual observations). To achieve this, the breeding and pre-breeding population data were partitioned into training and testing set in a 3:1 ratio with a stratification based on the target traits (chromameter b*value or dry matter content). The regression model developed in the training set was used to predict the trait values in the hold-out testing set. All model training and cross-validation analyses were implemented in the R caret package (Kuhn, 2008).

## 3.0 Results

### 3.1 Phenotypic variation for root yellowness parameters and dry matter content

Out of the evaluated clones, 81% of the breeding population and 52% of the pre-breeding population had white storage roots, while the remaining showed a range of yellow color (visual score of between 2 and 5), suggesting varying levels of carotenoid content (Figure 2). The average visual score of root yellowness was 1.30 (sd = 0.72) in the breeding population and 1.74 (sd = 0.95) in the pre-breeding population. The chromameter b* values showed a bi-modal distribution in the two populations (Figure 2). The first peak (b*values from 11 to 22) is associated with the white clones, while the second peak (b* values from 22 to 50) is associ-ated with the variations among the yellow clones. The average chromameter measures of yellow color intensity were 21.0 (sd = 6.12) and 26.2 (sd = 8.82) for breeding and pre-breeding populations, respectively. The dry matter content of the clones evaluated in the two populations was normally distributed (Figure 2), ranging from 11.2 to 47.4, with averages of 31.5 (sd = 5.92) in the pre-breeding population and 35.1 (sd = 4.80) in the breeding population.

**Figure 2:**
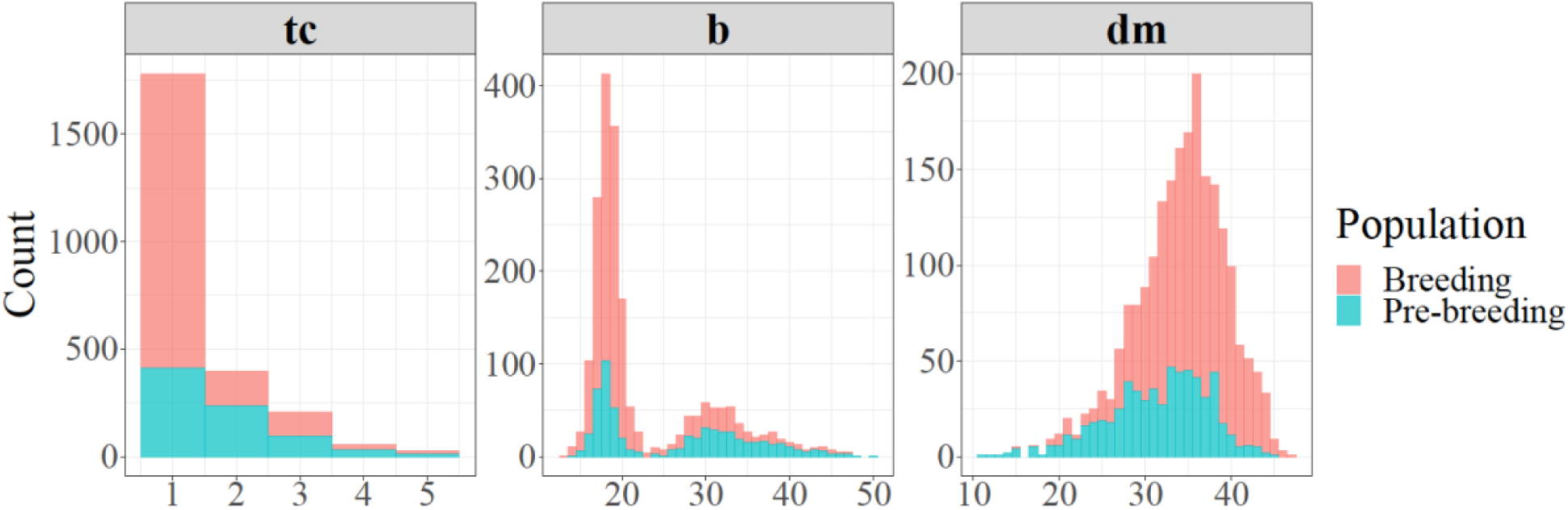
Frequency distribution of cassava genotypes for root yellowness intensity (color-chart (tc) and chromameter (b)) and dry matter content (dm %) in the breeding and pre-breeding populations

The broad-sense heritability of the visual assessment from the color chart and chromameter values were 0.87 and 0.88, respectively for the breeding population, and 0.81 and 0.93, respectively for the pre-breeding population (Table 2). The heritability estimate for dry matter content in the pre-breeding population (0.45) was low compared to that of the breeding population (0.70) (Table 2), indicating low genetic variation for the trait in the pre-breeding population.

**Table 2:** Broad-sense heritability calculated on a mean plot basis for root visual assessment, chromameter value, and dry matter content in the two populations

| Traits | Breeding population |  |  | Pre-breeding population |  |  |
| --- | --- | --- | --- | --- | --- | --- |
| | $\sigma^2_g$ | $\sigma^2_e$ | $H^2$ | $\sigma^2_g$ | $\sigma^2_e$ | $H^2$ |
| Visual assessment | 0.500 | 0.072 | 0.87 | 0.637 | 0.145 | 0.81 |
| Chromameter<br>b* value | 34.641 | 4.572 | 0.88 | 65.676 | 4.879 | 0.93 |
| Dry matter<br>content | 16.231 | 6.831 | 0.70 | 15.930 | 19.776 | 0.45 |

$\sigma_g^2$ is the clonal genotypic variance, $\sigma_e^2$ is the residual variance, and $H^2$ is the broad-sense heritability

The two measures of root yellowness intensity; visual assessment and chromameter b* value were significantly and positively correlated (~0.90) in the two populations suggesting that visual scoring is a good proxy for yellow-color intensity. Significant negative correlations ranging from −0.29 to −0.23 were observed between root yellowness and dry matter content in the two populations. However, a lower magnitude of correlation coefficient was observed between visual assessment and dry matter content (−0.23) as well as between chromameter b* value and dry matter content (−0.24) in the pre-breeding population.

### 3.2 Technical validation of Kompetitive Allele-Specific PCR (KASP) assays

#### 3.2.1 SNP call rate, call clarity, and genotypic frequencies

All markers were successfully converted to allele-specific KASP assay. The call rate and clarity were high for a wide range of DNA dilution levels tested during marker development indicating that the assays are robust and suitable for routine use (Supplementary Figure 2). The overall call rate was above 98 % for all the markers in the two populations genotyped (mean = 99%, sd = 0.53) (Supplementary Table 1). As expected, three distinct clusters were observed for all the SNPs except for marker S5_3387558 where the frequency of cluster TT was very low (Supplementary Figure 3).

Allelic and genotypic frequencies of the markers are presented in Supplementary Figures 4 and 5, respectively. The favorable alleles across all the carotenoid-linked markers were more common in the pre-breeding population (range 11% to 34%) compared to the breeding population (range 3% to 11%) (Supplementary Figure 4). The favorable allele A at marker S1_24155522 had a frequency of 34% and 11% in the pre-breeding and breeding population, respectively (Supplementary Figure 4). More than 15% of the individuals were homozygous for allele A at this marker in the pre-breeding population (Supplementary Figure 5). The percentage was much lower in the breeding population with only 2.3% of the individuals fixed for the same allele. In the two populations, between 0.4% to 7.3% of the individuals were fixed for the favorable alleles at the three remaining markers suggesting an opportunity to use these markers to increase their frequencies in the population (Supplementary Figure 5).

For dry matter content, the favorable alleles at the linked SNPs occurred at intermediate to high frequencies ranging from 28% to 76% in both populations (Supplementary Figure 4). The percentage of individuals that were fixed for the favorable alleles was higher in the breeding than the pre-breeding population for this trait (Supplementary Figure 5). About 27% to 53% of the individuals in the pre-breeding population were fixed for the unfavorable alleles (Supplementary Figure 5).

#### 3.2.2 Biological validation

##### 3.2.2.1 Allelic substitution effects on carotenoid and dry matter contents

Significant pairwise differences between genotypic classes at all the markers associated with carotenoid content were observed (Figure 3). Most of the markers displayed additive mode of action with individuals carrying two copies of the favorable alleles having a higher intensity of root yellowness (b*) while those that are fixed for non-favorable alleles had white roots. For instance, the mean b* values for genotype classes AA, CA, and CC for marker S1_24155522 were 38.53 ± 2.85, 31.64 ± 3.89, and 18.37 ± 2.48, respectively in the pre-breeding population (Figure 3b).

**Figure 3:**
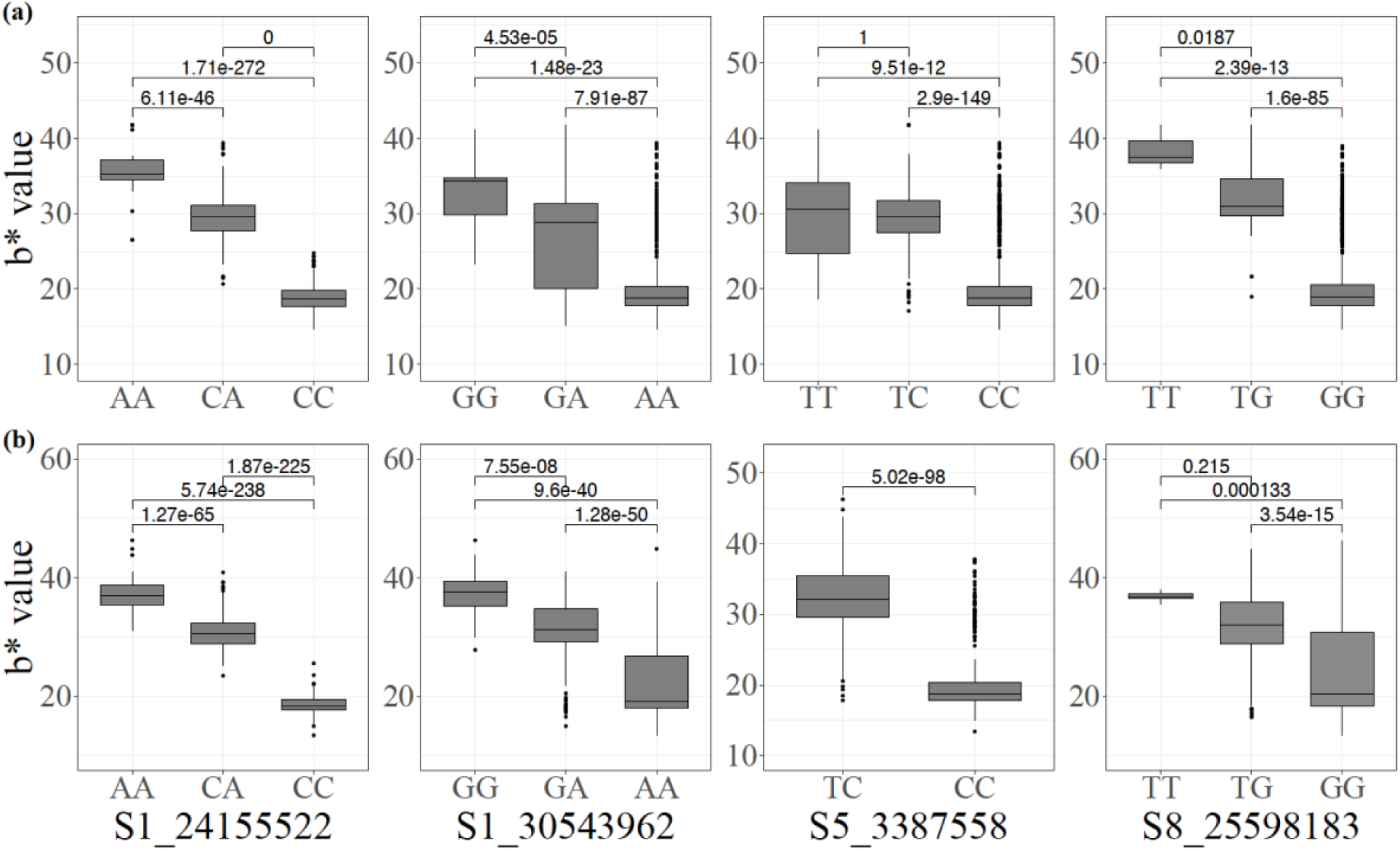
Allelic substitution effects of the markers associated with increased carotenoid content in the (a) breeding, and (b) pre-breeding populations (For marker S5_3387558, the mean and standard deviation cannot be estimated because one genotype had TT)

The genotype classes at the dry matter content-linked markers were not as differentiated as those for carotenoid content (Figure 4). Nonetheless, significant differences were observed among the genotypes at marker S6_20589894 in the two populations. In the pre-breeding population, there was no significant difference among CC, CT, and TT at marker S12_5524524 (Figure 4).

**Figure 4:**
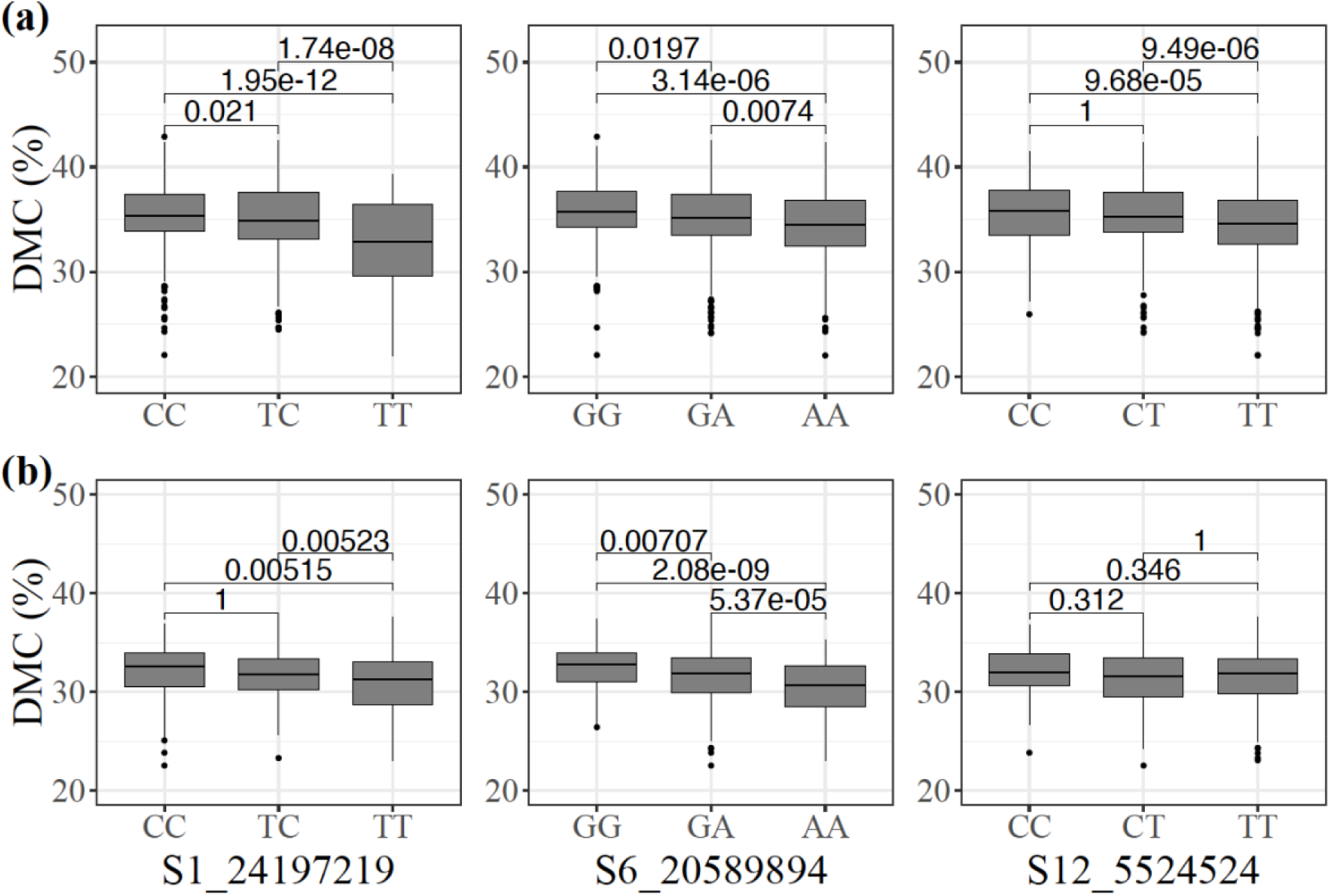
Allelic substitution effects of the markers associated with increased dry matter content (DMC) in the (a) breeding, and (b) pre-breeding populations

#### 3.2.2.2 Marker-trait regression, confidence intervals, and models’ predictive performances

The estimates of marker-trait regression parameters from bootstrap resampling analysis for the two traits are presented in Figures 5 and 6. The regression model with all the four markers for carotenoid variation produced R^2^ values of 0.85 in the breeding and 0.91 in the pre-breeding population. SNP S1_2415552 had the strongest effect on variation in root yellowness. The effect size of having a single copy of a favorable allele (A) on the increase in root yellowness intensity (chromameter b* value) was 10.8 and 12.1 in the breeding and pre-breeding populations, respectively. Having two copies of the same allele resulted in an even larger effect size of 15.5 and 17.8, respectively, in the two populations. The confidence intervals of these marker genotypes were narrow, indicating higher precision of the marker prediction. After controlling for the major locus (S1_24155522), the other three markers had a low to moderate effect on the trait (Supplementary Figure 6). The effect sizes of the minor SNPs were more significant in the breeding compared to the pre-breeding population, particularly for markers S5_3387758 and S8_25598183.

**Figure 5:**
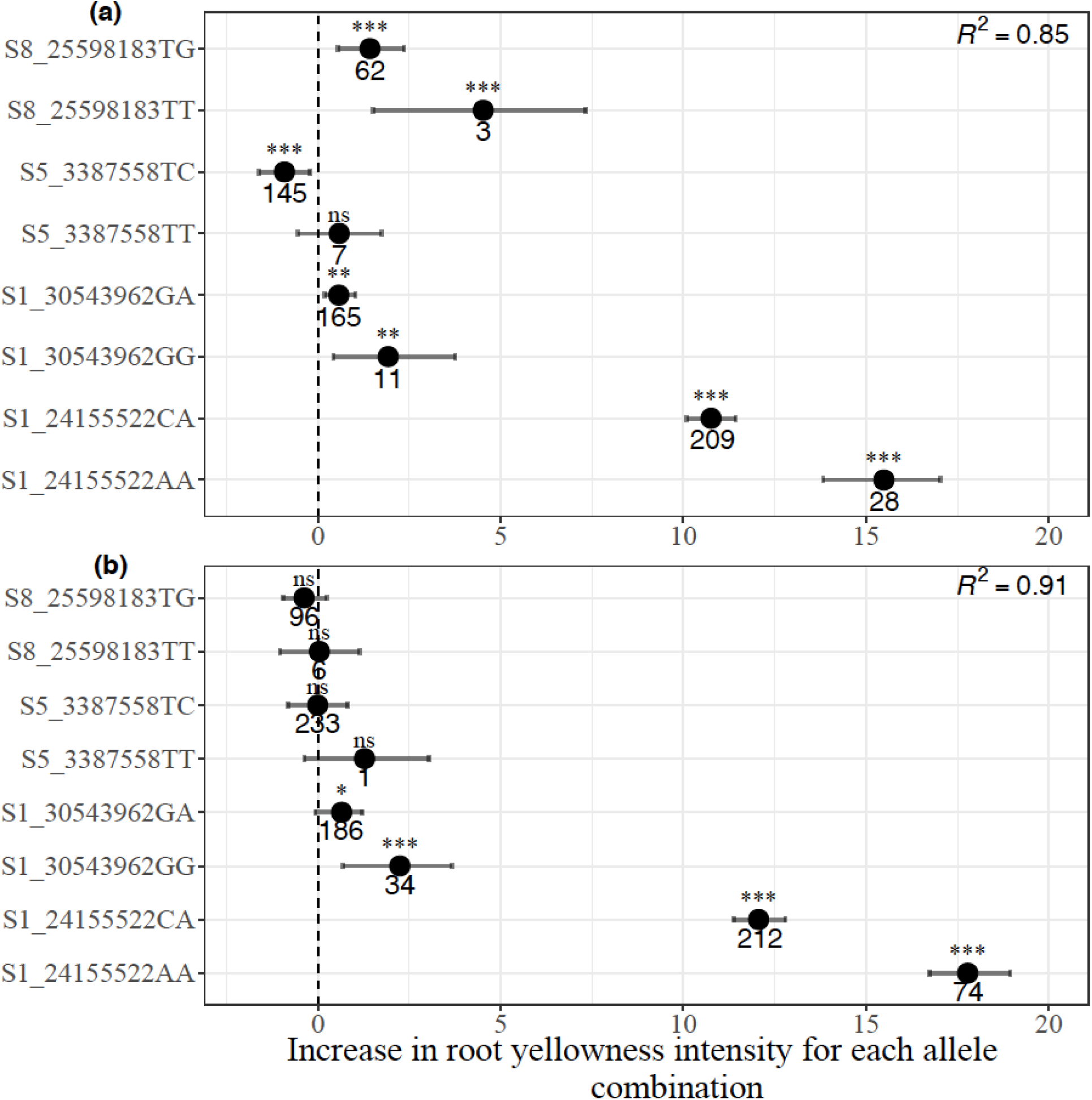
Distribution of marker allelic effects associated with increased carotenoid content in (a) breeding, and (b) pre-breeding populations

**Figure 6:**
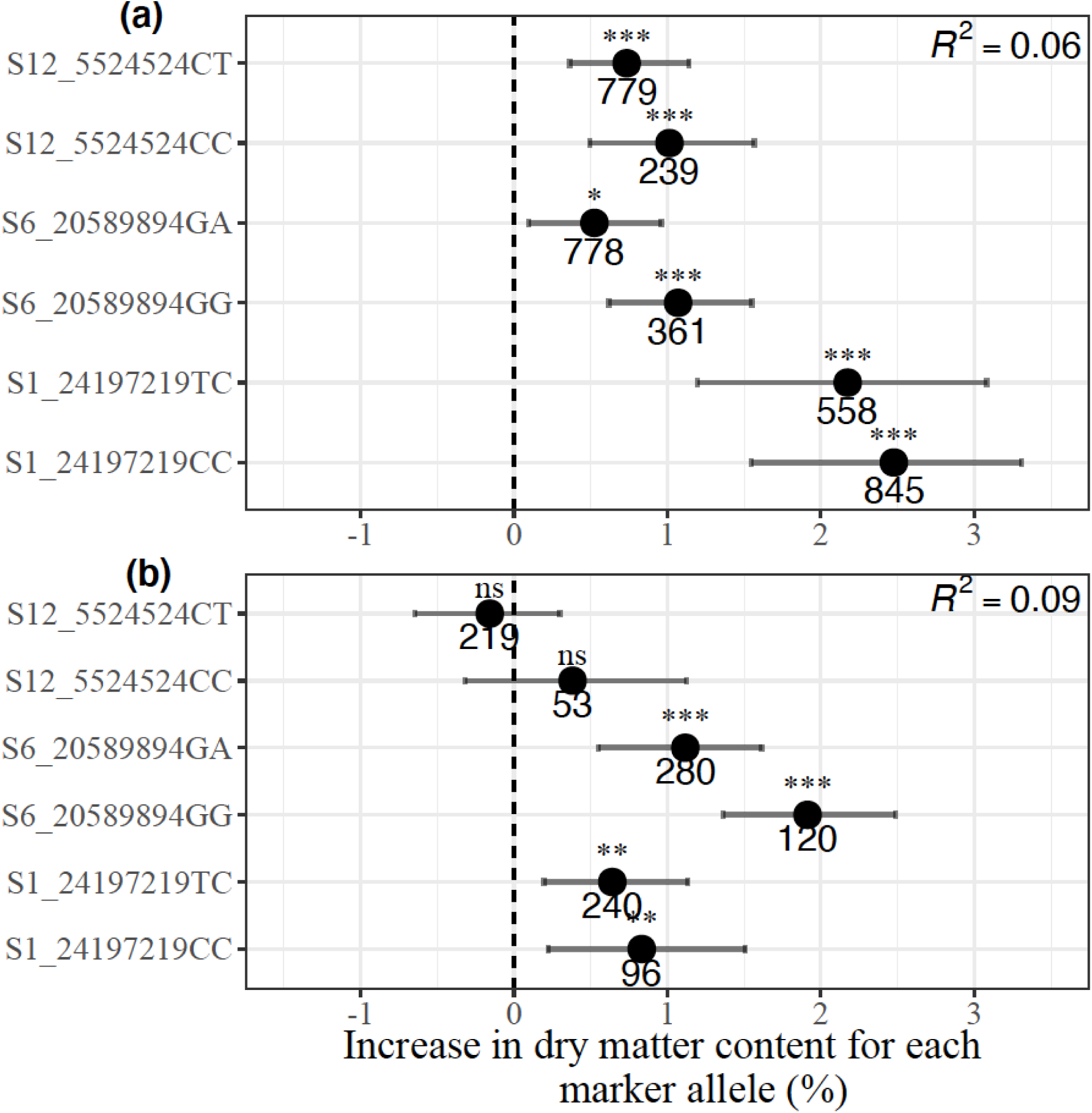
Distribution of the marker allelic effects associated with increased dry matter content in (a) breeding, and (b) pre-breeding populations

The regression model with all three markers for dry matter content produced low R^2^ values of 0.06 in the breeding and 0.09 in the pre-breeding population. Having two copies of favorable alleles across all SNPs was associated with an increase in dry matter content percentage from between 1.01 and 2.50 percentage units in the breeding population. A similar direction of effects was observed in the pre-breeding population except for marker S12_5524524 which did not contribute to the multiple regression model. A notable observation is a reversal in the effects of markers S1_24197219 and S6_20589894 across the two populations, suggesting a QTL by genetic background interaction.

The predictive accuracy of the carotenoid markers from the cross-validation regression analysis ranged from 0.84 to 0.91 with a mean of 0.87. In the pre-breeding population, the value was higher and approximately 0.90 in the training and testing sets (Table 3, Supplementary Figure 7). However, low predictive accuracy values were obtained for dry matter content-linked markers in the breeding population (0.06 for the training set and 0.05 for the testing set) and pre-breeding population (0.06 for the training set and 0.15 for the testing set) (Table 3, Supplementary Figure 7). In the breeding population, RMSE and MAE values for carotenoid markers were 1.88 and 1.43, respectively in the training set and 2.03 and 1.52, respectively in the testing set (Table 3). The values of RMSE and MAE were 2.31 and 1.71, respectively in the training set, and 2.35 and 1.68 in the testing set of the pre-breeding population. These values were slightly higher for dry matter content-markers in both populations compared to that of carotenoid content-markers. The use of RMSE and MAE is very common in model evaluation, and they are good measures of prediction accuracy.

**Table 3:** Prediction performance metrics of the markers associated with increased carotenoid and dry matter contents in the training and testing sets of the breeding and pre-breeding populations

| Traits | Populations |  | N | R<sup>2</sup> | RMSE | MAE |
| --- | --- | --- | --- | --- | --- | --- |
| Chroma meter b* value (Carotenoid content) | Breeding | Training set | 1030 | 0.84 | 1.88 | 1.43 |
|  |  | Testing set | 345 | 0.84 | 2.03 | 1.52 |
|  | Pre-breeding | Training set | 396 | 0.91 | 2.31 | 1.71 |
|  |  | Testing set | 133 | 0.90 | 2.35 | 1.68 |
| Dry matter content (%) | Breeding | Training set | 1102 | 0.06 | 3.14 | 2.49 |
|  |  | Testing set | 368 | 0.05 | 3.20 | 2.48 |
|  | Pre-breeding | Training set | 402 | 0.06 | 2.59 | 2.10 |
|  |  | Testing set | 136 | 0.15 | 2.56 | 2.10 |

N=Number of observations, R<sup>2</sup>= Prediction accuracy, RMSE = Root mean square error, MAE = Mean absolute error

## 4.0 Discussion

The present study focused on the development and validation of markers for carotenoids and dry matter contents: two traits that are of primary importance to cassava breeding programs worldwide (Sánchez et al., 2006; Okechukwu and Dixon, 2008; Bouis et al., 2011; Saltzman et al., 2013; Talsma et al., 2013). As part of breeders’ toolbox for MAS, such markers are expected to address the challenges associated with vitamin A deficiency (VAD) and higher demand for varieties with higher dry matter content. Vitamin A deficiency is a widespread nutritional public health problem in sub-Saharan Africa, with women and children being the most affected (Gegios et al., 2010; Stephenson et al., 2010). Breeding of clones with enhanced carotenoid levels is one of the most cost-effective and sustainable approaches to help the communities burdened by VAD (Pfeiffer and McClafferty, 2007; Bouis et al., 2011; Talsma et al., 2013).

While we have explored the performance of the markers in the IITA pre-breeding and breeding populations, these assays should have wide application in other breeding programs where the QTLs are present and are linked to the same SNP alleles. More importantly, these markers can be used for rapid mobilization of the favorable allele in new populations if used with parents that are known to carry the associated trait alleles.

Trait discovery in cassava has been an active area of research with the advent of genome-wide SNP markers from genotyping-by-sequencing (Esuma et al., 2016; Wolfe et al., 2016; Rabbi et al., 2017, 2020; Udoh et al., 2017; Ikeogu et al., 2019). However, these trait discoveries have not been translated to deployable assays, obscuring their utility in MAS. Here, we have provided a framework for translating the outputs from genetic mapping to a set of easy-to-use, robust and predictive allele-specific uniplex assays. The framework includes both technical and biological validation of the assays in a range of diverse germplasm to ascertain the relevance of the markers for predicting the trait values in independent populations. The KASP SNP platform was chosen due to its amenability for genotyping of any combination of individual samples and maker assays, and ease of automation to achieve high-throughput population screening (Semagn et al., 2014; Ige et al., 2021). The designed SNP assays were found to work under a wide range of DNA concentrations. Even though the tightness of the cluster plots differed between the standard and low DNA concentrations, they were sufficiently distinct to allow for high genotype call rate and call clarity. This suggests that the assays are expected to work under diverse DNA concentrations and most likely from different sample preparation methods including fresh, frozen, lyophilized, or oven-dried (Semagn et al., 2014).

The best way to measure the predictive ability of a model is to test it on a dataset that is independent of the data used to train the model (Wani et al., 2018). The *k*-fold cross-validation, where the original dataset is randomly partitioned into equally sized *k*-subsets (a single subset is retained as the validation data for testing the model, and the remaining *k* - 1 subsets are used as training data) is one of the most commonly used cross-validation methods (Refaeilzadeh et al., 2009; Mathew et al., 2015). It is routinely used to assess genomic prediction accuracies (Okeke et al., 2017; Andrade et al., 2019; Phumichai et al., 2022). To our knowledge, this is the first study to use this metric for marker validation in cassava. In the present study, the performance of the regression model in an independent data set, that is, the testing set in terms of predictive accuracy for chromameter b* values were 0.84 in the breeding population and 0.90 in the pre-breeding population. These values are quite similar to what was obtained in the training sets suggesting that the models developed are stable and reliable. The low values of RMSE and MAE recorded in the breeding population compared to that of the pre-breeding population indicated that the markers are more accurate in predicting the carotenoid content in the breeding population. Both measures of cross-validation accuracy for this trait suggest that the designed assays can be deployed for routine use in breeding pipelines with carotenoid biofortification as a breeding goal. On the other hand, the predictive accuracy of the dry matter content markers (mean = 0.08) across populations was not as high as the values obtained for carotenoid content markers. This could be due to the quantitative nature of the trait (Kawano et al., 1987) and is generally characterized by moderate heritability values as reported in this study. In the discovery population, Rabbi et al. (2020) also reported low predictive ability (R^2^ < 0.11) of these markers.

Moreover, for both traits, we used a bootstrapping regression approach to provide robust estimates of allele substitution effect and their confidence intervals through repeated resampling from the data (Fox and Weisberg, 2018). The multiple regression analysis of carotenoid content markers revealed that marker S1_24155522 was the main driver in carotenoid accumulation while the other markers played additional but minor roles. This result is consistent with earlier observations that the PSY2 gene which hosts marker S1_24155522 is a key rate-limiting step in the carotenoid pathway in cassava (Welsch et al., 2010; Rabbi et al., 2020). In a candidate gene-based association study, Udoh *et al*. (2017) reported total carotenoids content and β-carotene were significantly associated with this marker which occurs at position 572 of the PSY2 gene. Indeed, the previously identified SNPs from other candidate genes such as *lcyE, lcyB* and *crtRB* were hardly significantly associated with the trait (Udoh et al., 2017). On the other hand, markers S1_24197219 and S6_20589894 had small but significant effects on dry matter content in both populations while marker S12_5524524 showed an effect in the pre-breeding population. Marker S6_20589894 was reported to occur in close proximity to gene Manes.06G103600 (Bidirectional sugar transporter Sweet4-Related) which mediates fructose transport across the tonoplast of roots (Rabbi et al., 2020).

While we have assessed the performance of selected markers across the two diverse populations, we acknowledge that these markers may be tagging only a subset of major loci underlying the studied traits, particularly dry matter content. Ongoing and future GWAS and biparental QTL mapping studies will likely uncover additional QTLs. Such markers can be validated using the framework provided in this study and incorporated into the breeders’ toolset, thus increasing the accuracy of predicting these traits. Moreover, other traits that are of importance for which major associations have recently been reported but not converted to marker assays include cyanogenic potential (Ogbonna et al., 2020), cassava green mite (Rabbi et al., 2020), cassava brown streak disease (Kayondo et al., 2018) and root mealiness (Uchendu et al., 2021). A major caveat of our study is the use of single-marker assays to tag each major locus for the two traits. The top SNPs at these loci are expected to be tightly linked to the causal allele, based on the large GWAS population used in the discovery with more than 5000 individuals genotyped at more than 100K genome-wide positions. However, factors such as independent emergence or evolution of favorable alleles at specific genes and nearby SNP can result in non-perfect association, hence resulting in false-positive and false-negative. This and other limitations of single marker MAS can be addressed by a haplotype-based approach through for example amplicon sequencing (AmpSeq) of targeted genomic regions (Yang et al., 2016). Further work is required to establish the viability of Amplicon Sequencing as a platform for haplotype-based MAS in cassava.

## 5.0 Conclusion

We have successfully developed and validated seven stable and novel carotenoid content and dry matter content markers in two independent cassava populations. A standardized schema presented in this study can be used for future cassava marker validation processes. All the markers validated had a high call rate (>98%) and clear allele calls using the KASP platform. Marker S1_24155522 located within the PSY2 gene had the highest effects on carotenoid content compared to small other markers because of the role of the gene in the carotenoid pathway in cassava roots. While the markers for carotenoid content can be used to select “winning genotypes” because of their high predictive power (R^2^ > 0.80), those for dry matter content are more useful for eliminating individuals that are homozygous for the non-favorable alleles rather than identifying those with high dry matter content levels (R^2^ < 0.10).

## Data Availability Statement

Data are available at cassavabase.org

## Author Contributions

Designed study: ADI, IYR and PK

Developed the pre-breeding population: SM-W, JN

Performed the experiment: ADI, RU,

Analyzed the data: ADI, GJB, and IYR

Drafted the manuscript: ADI

Revised the manuscript: BO, GJB, ISK, EGNM, CE, EYP, PK, HC, and IYR

All authors have read, edited, and approved the current version of the manuscript.

