## Supplementary Files for "Validation of SNP markers for marker-assisted selection of genotypes with increased carotenoid and dry matter contents in cassava"

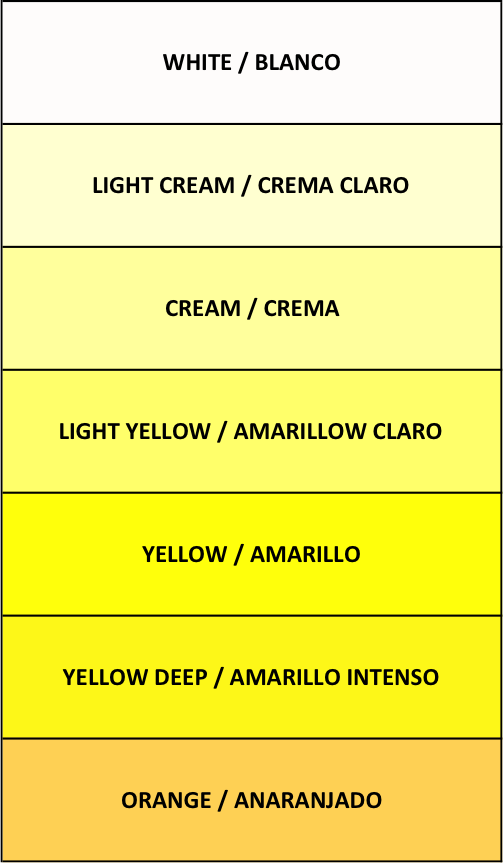

Supplementary Figure 1: Standard color-chart for scoring cassava tissue for carotenoid content

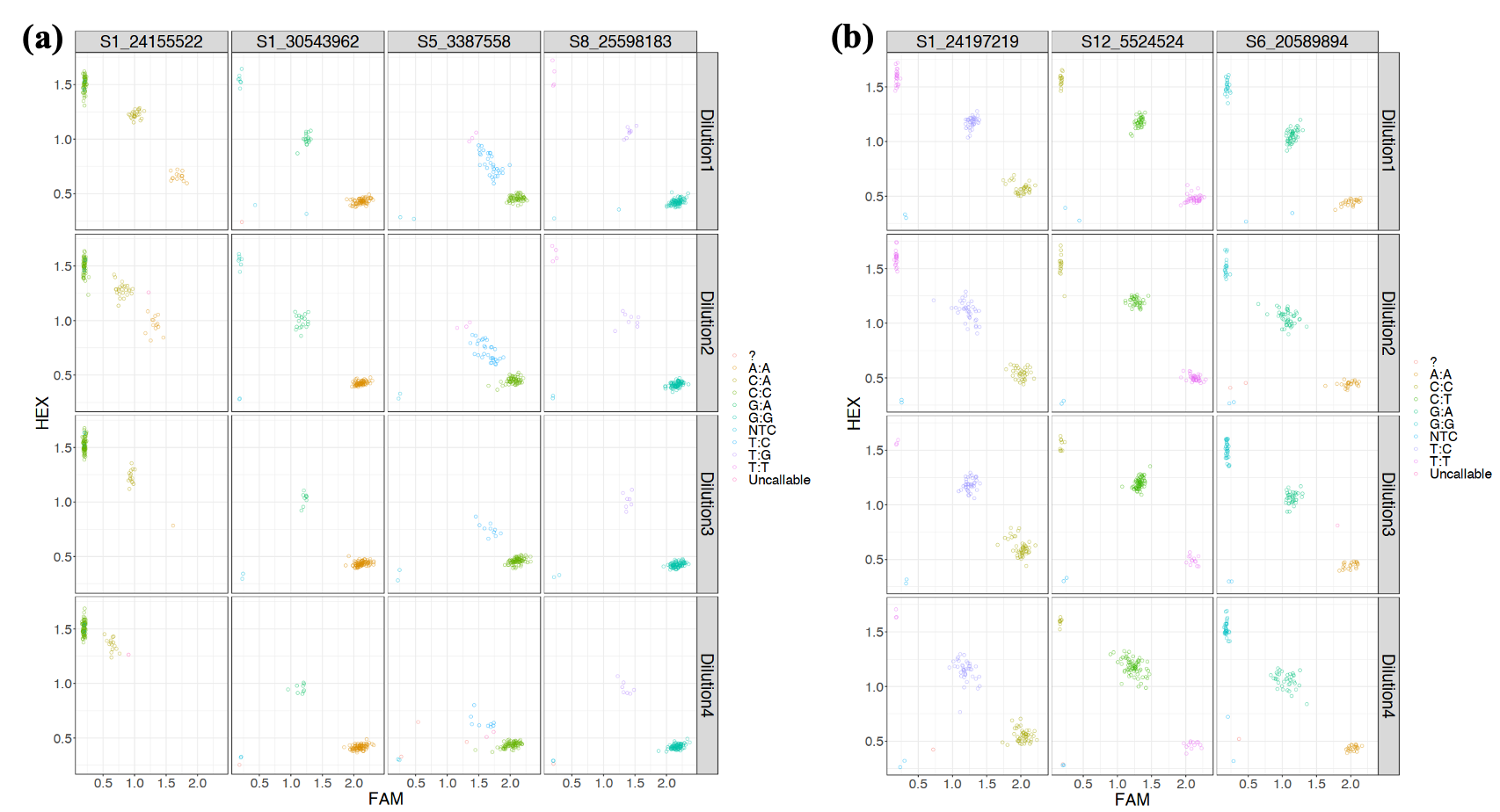

Supplementary Figure 2: Scatter plots of genotypic classes from the KASP marker assays associated with increased (a) carotenoid content and (b) dry matter content in the 188 diverse cassava germplasm used for the technical validation at four DNA concentrations (Dilution 1= 10X, Dilution 2 = 100X, Dilution 3 = 24X, Dilution 4 = 240X)

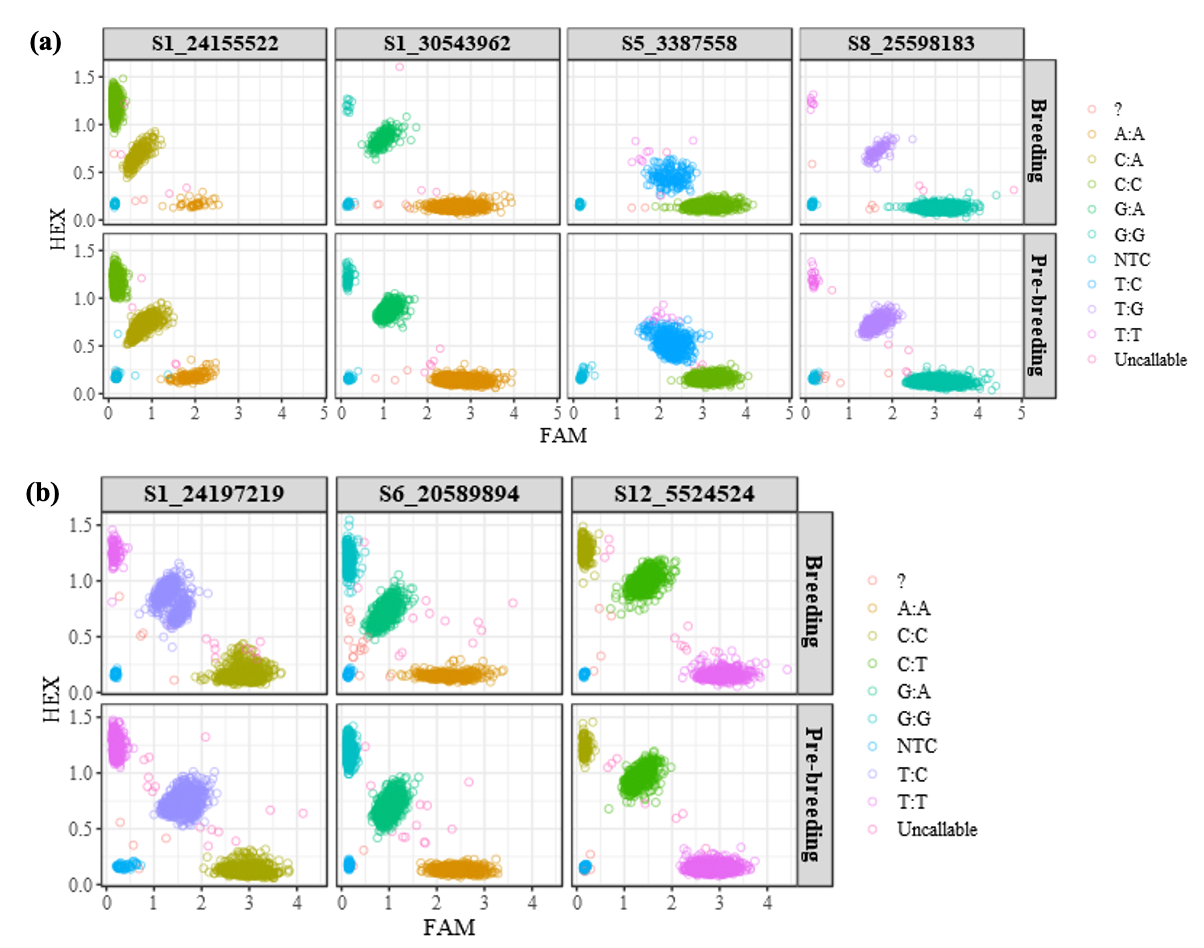
Supplementary Figure 3: Scatter plots of genotypic classes from the KASP marker assays associated with increased (a) carotenoid content and (b) dry matter content in the breeding and pre-breeding populations

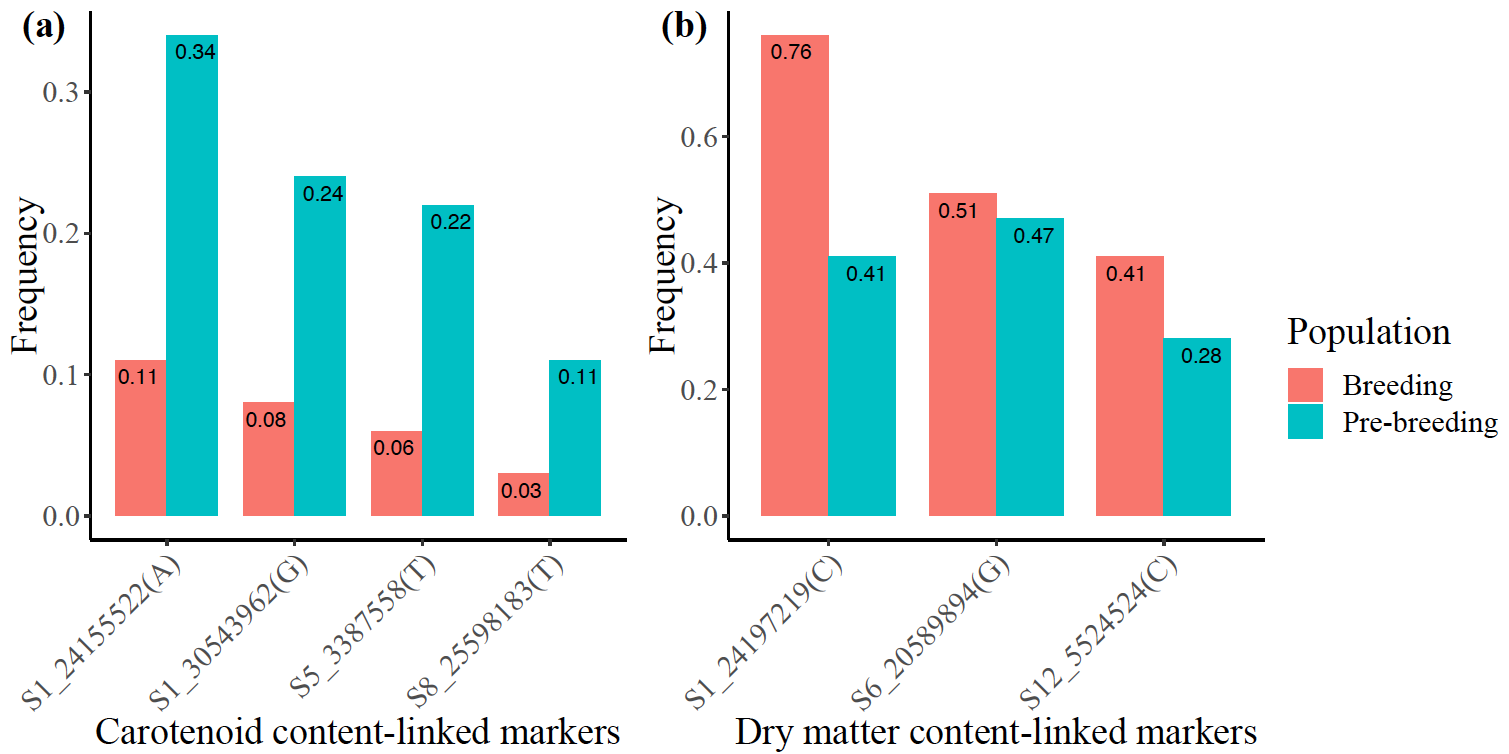
Supplementary Figure 4: Frequencies of favorable alleles at the markers associated with increased (a) carotenoid content and (b) dry matter content in the breeding and pre-breeding populations

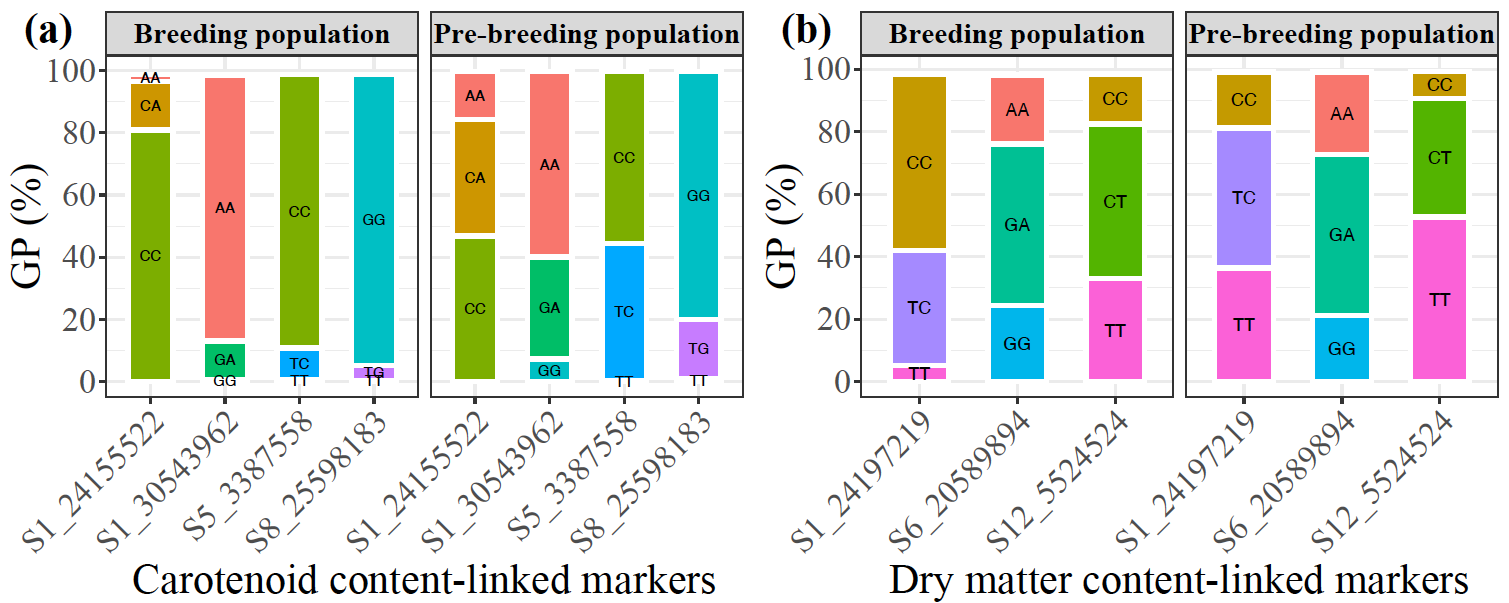

Supplementary Figure 5: Marker genotype frequencies associated with increased (a) carotenoid content and (b) dry matter content in the breeding and pre-breeding populations

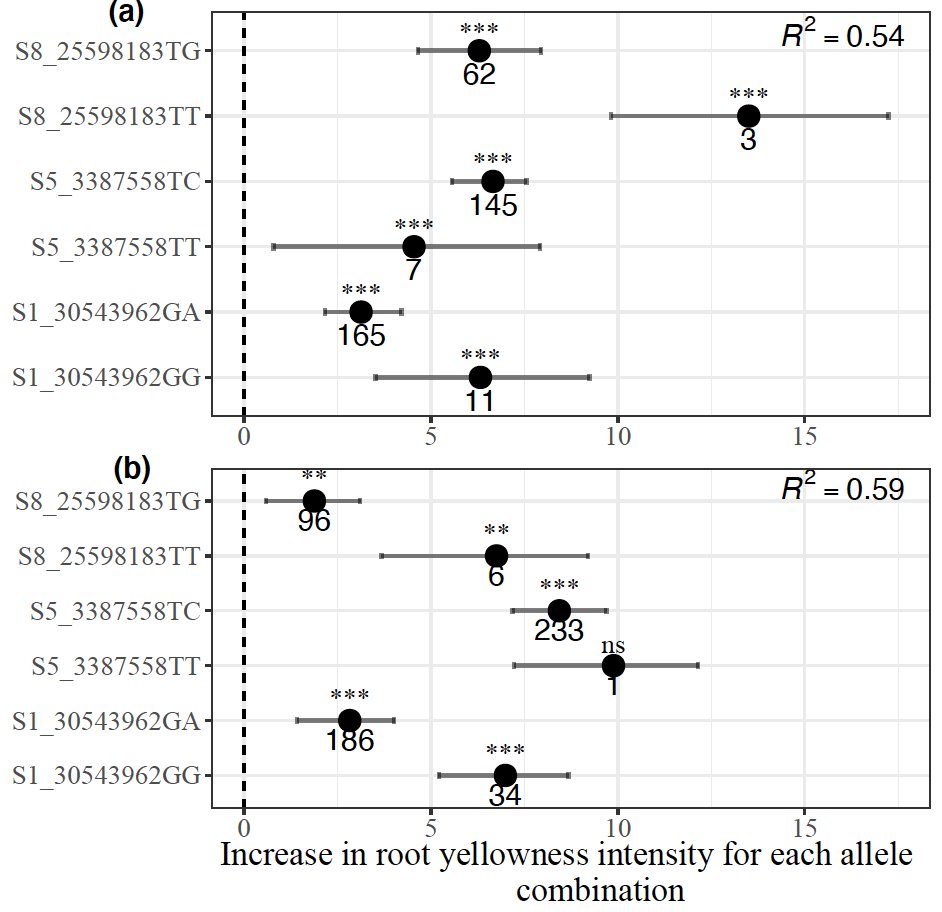

Supplementary Figure 6: Distribution of marker allelic effects associated with increased carotenoid content in (a) breeding, and (b) pre-breeding populations (The multiple linear regression model excluded marker S1_24155522)

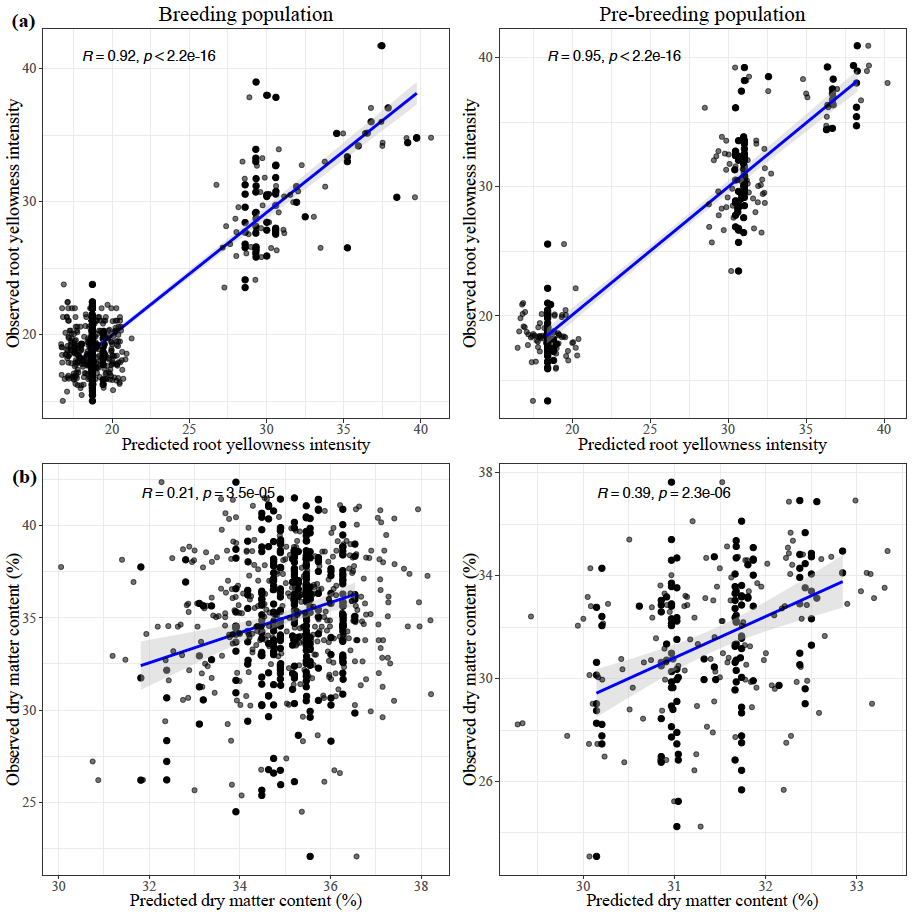

Supplementary Figure 7: Predictive accuracies (r) of the markers associated with increased (a) carotenoid content and, (b) dry matter content for the testing sets of the breeding and pre-breeding populations
